# DNA shape complements sequence-based representations of transcription factor binding sites

**DOI:** 10.1101/666735

**Authors:** Peter DeFord, James Taylor

## Abstract

The position weight matrix (PWM) has long been a useful tool for describing variation in the composition of regions of DNA such as transcription factor (TF) binding sites. It is difficult, however, to relate the sequence-based representation of a DNA motif to the biological features of the interaction of a TF with its binding site. Here we present an alternative strategy for representing DNA motifs – called Structural Motif (StruM) – that can easily represent different sets of structural features. Structural features are inferred from dinucleotide properties listed in the Dinucleotide Property Database. StruMs are able to specifically model TF binding sites, using an encoding strategy that is distinct from sequence-based models. This difference in encoding strategies makes StruMs complementary to sequence-based methods of TF binding site identification.

## 1 INTRODUCTION

The human body has trillions of cells and hundreds of cell types that have many distinct morphologies and functional roles [1, 2]. Yet each of these cells has an identical copy of the underlying developmental program encoded in the DNA [3, 4]. The diversity of form and function that is observed is only possible through tight control of the expression of genes throughout the developmental process [5, 6, 7]. Understanding the mechanisms guiding this control will provide insight into the development of complex multicellular organisms, and associated diseases.

Transcription is controlled by the activity of *cis*-regulatory modules; regions of DNA that, when bound by the appropriate factors, serve to promote or inhibit transcription, and can exist in many different states of activity [7, 8]. The best computational approaches to identifying these modules rely on large amounts of experimental data, primarily targeting transcription factor binding (e.g. [9]). However, the available data is quite limited, with only a handful of experiments available at most for many cell types. Experimentally generating a comprehensive dataset of protein-DNA interactions for all cell types at this point is intractable due to limitations of the number of cells required for some experiments, the quality of reagents, and cost.

Computational methods that predict transcription factor (TF) binding complement the available data, and allow for extrapolation across rare or otherwise unwieldy cell types. Historically this is done by representing the chains of nucleotides that make up DNA as sequences composed of an alphabet of 4 letters. Patterns within these sequences can be identified, for example using letter frequencies at each position within a set of aligned binding sites as in the position weight matrix (PWM) [10]. This can be quite effective, and has been shown to perform well for many transcription factors [11]. Some transcription factors have been shown to be especially well represented in this manner, displaying extreme sequences preferences enforced through base pair-specific contacts along the major groove. This mode of binding site recognition by transcription factors is known as direct- or sequence-readout.

In reality, DNA is a complex three-dimensional macromolecule that is tightly packed into the nucleus. Other transcription factors have been shown to take advantage of the three dimensional shape of DNA molecule to recognize their binding sites in a mode known as indirect- or shape-readout [12, 13, 14]. For example, it has been shown that the narrowing of the minor groove and corresponding increase in electrostatic potential drives interactions with positively charged arginine side chains in Oct-1/PORE complex binding [12].

The sequence representation of DNA and the associated TF binding site (TFBS) models are in reality an abstraction of the chemical and physical interactions of the protein molecules with the DNA. As indicated above, even sequence-readout relies on the proper 3D positioning and electrostatic compatibility of hydrogen bond donors and acceptors between the two molecules.

We hypothesized that TF binding preferences could be modeled using estimates of DNA shape parameters across the binding site, and this would provide increased discrimination over sequence-based approaches. To this end we designed a set of methods adapting the time-tested position weight matrix to incorporate DNA shape instead of sequence, known as Structural Motifs (StruMs). StruMs specifically model TFBSs and are complementary to sequence-based methods.

## 2 MATERIALS AND METHODS

### 2.1 Structural Motifs

The StruM is an extension of the PWM [10, 15]. Each position-specific feature is assumed to be independent allowing for log-probabilities to be combined additively, and the model finds a simple distribution across each of these features. The construction of a motif requires two things: 1) aligned sequences corresponding to binding sites; and 2) a method for estimating shape features. It has been shown that, as with proteins, the primary sequence of the DNA plays a large role in determining the local shape of the DNA [12]. Here the Dinucleotide Property Database (DiProDB) [16] is used to estimate shape parameters for each dinucleotide in the sequence.

#### 2.1.1 Definition

Given a set of *n* training sequences, each sequence is converted to a structural representation. In this case each consecutive dinucleotide is looked up in DiProDB, and that column is appended to the feature vector. The length of this vector is *k* (the length of the binding site) times *p* (the number of shape features being considered); this value *k* · *p* is noted hereafter as *m*. This set of training structures (*D*) is used to compute the parameters (*ϕ*). These are represented as a mean (*µ*) and standard deviation (*σ*) for each feature at each position.

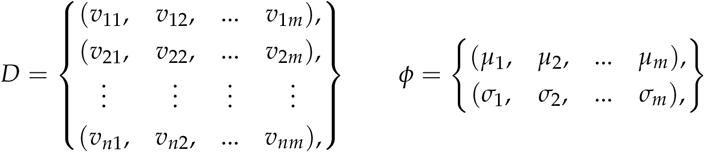

The model begins with the assumption that the DNA shapes preferred for interaction with a given transcription factor at any given position specific feature (*v*_*j*_) have an optimum shape, and sample adjacent shapes according to a normal distribution.

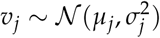

Assuming that each feature and each position is independent, then calculating the score (*s*) for the *i*-th sequence becomes:

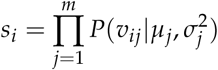

In order to avoid underflow issues during computation, all calculations are done in log space.

### 2.2 Data

All experimental data is obtained from ENCODE [17]. The ChIP-seq data used Transcription Factor ChIP-seq data from K562 cells mapped to *hg19*. This data was filtered for targets that were annotated as being sequence specific transcription factors in [18]. Specifically the conservative IDR thresholded peaks were downloaded in the ENCODE narrowPeak BED format. TF family assignments were done using the assignments in [19]. Accession numbers for all datasets used are available in the Supplementary Information.

### 2.3 ChIP peak classification

#### 2.3.1 *De novo* motif finding

For each TF analyzed, the sequences for the top 500 peaks were retrieved based on signal enrichment. The sequence corresponding to 100 bp around the peak identified in the BED file was then extracted. A PWM was learned using `meme-chip` [20] with the parameters `-norand -meme-nmotifs 1 -dreme-m 0 -spamo-skip -dna -nmeme 500 -seed`. The model was left as the default (`zoops`, and the random seed was set by hashing each TF’s accession number. The sites reported by MEME as being used to train the final motif were extracted from the output. These aligned binding sites were used to compute position specific frequencies for mononucleotides (PWM) [10] and dinucleotides (dinucleotide weight matrix, DWM) [21]. After translation into structural space, the distribution of shape values at each position was calculated for the StruM. This variation will be referred to as the Maximum Likelihood StruM (ML-StruM). Additionally, the 100 bp centered sequences were used to fit an additional structural motif by expectation-maximization, matching its length to that of the PWM learned by MEME (StruM). Similarly to the incorporation of pseudocounts in nucleotide frequency estimations, a minimum threshold of 0.1 was imposed on *σ* for the StruM.

#### 2.3.2 Motif performance

For the next 500 sequences in the ChIP-seq experiment, the maximum score for each motif type was calculated. To generate scores for a matched set of negative sequences, two strategies were employed: shuffling these testing sequences, or taking 100 bp flanking sequences from the top 500 peaks. Upon generation of the negative set, the scoring process was repeated. A simple threshold was varied across the scores to generate a receiver operating characteristic (ROC) curve and a precision-recall curve (PRC) and the area under the curve (AUC) was calculated. This process was repeated for 355 TF ChIP-seq experiments in K562 cells.

#### 2.3.3 Specificity of StruM by TF family

Using TF family assignments in TFClass ([19]), the second 500 sequences for each TF was pooled by TF family. For each TF, the second 500 sequences for that TF were compared to 500 sequences randomly sampled from the other TF families. As a control, 500 sequences randomly selected from that TF’s family pool were compared to the subset from the other families.

#### 2.3.4 Complementarity of methods

To assess the complementarity of StruMs with sequence-based methods, the scores for each sequence were passed as a vector of length two (PWM score, StruM score) to a logistic regression classifier as implemented in Python’s `sklearn` [22, 23]. Ten-fold cross validation was performed with `sklearn.cross_validation. cross_val_score`, and the average AUC was retrieved.

### 2.4 Proximity of PWMs and StruMs

Given the motif derived by MEME for the ChIP-seq experiments described above, FIMO was used to scan the input sequences for statistically significant matches, as executed by `meme-chip` [24, 20, 25]. These significant matches were merged if they were within 100 bp of each other.

The sequences within 100 bp of the center of the clusters of significant matches were extracted, and scored with the EM derived StruM. The highest scoring position for each sequence was recorded. The average distance between the genomic locations of these best StruM matches to the nearest significant match to the PWM was then computed.

### 2.5 Program Versions

All programs and packages used in this analysis were downloaded and installed on a system running Ubuntu 16.04.6 LTS (GNU/Linux 4.4.0-150-generic x86_64) using Conda (4.6.2). The following packages and versions were used: MEME-suite (meme-chip, fimo) (4.12.0), Bedtools (2.27.1), Sci-kit learn (0.20.1), numpy (1.15.4), matplotlib (2.2.3), python (2.7.15), scipy (1.1.0).

## 3 RESULTS

### 3.1 Overview of the StruM model

The traditional representation of binding preferences for transcription factors, or their motifs, is the PWM. Given a set of sequences that are known to be bound by a TF, e.g. GATA1 (Figure 1a), these sequences can be aligned by the binding site (Figure 1c). Assuming that each of the positions in the binding site are independent, they can each be represented by a distribution of nucleotide frequencies; one distribution per position in the motif (Figure 1b).

**Figure 1:**
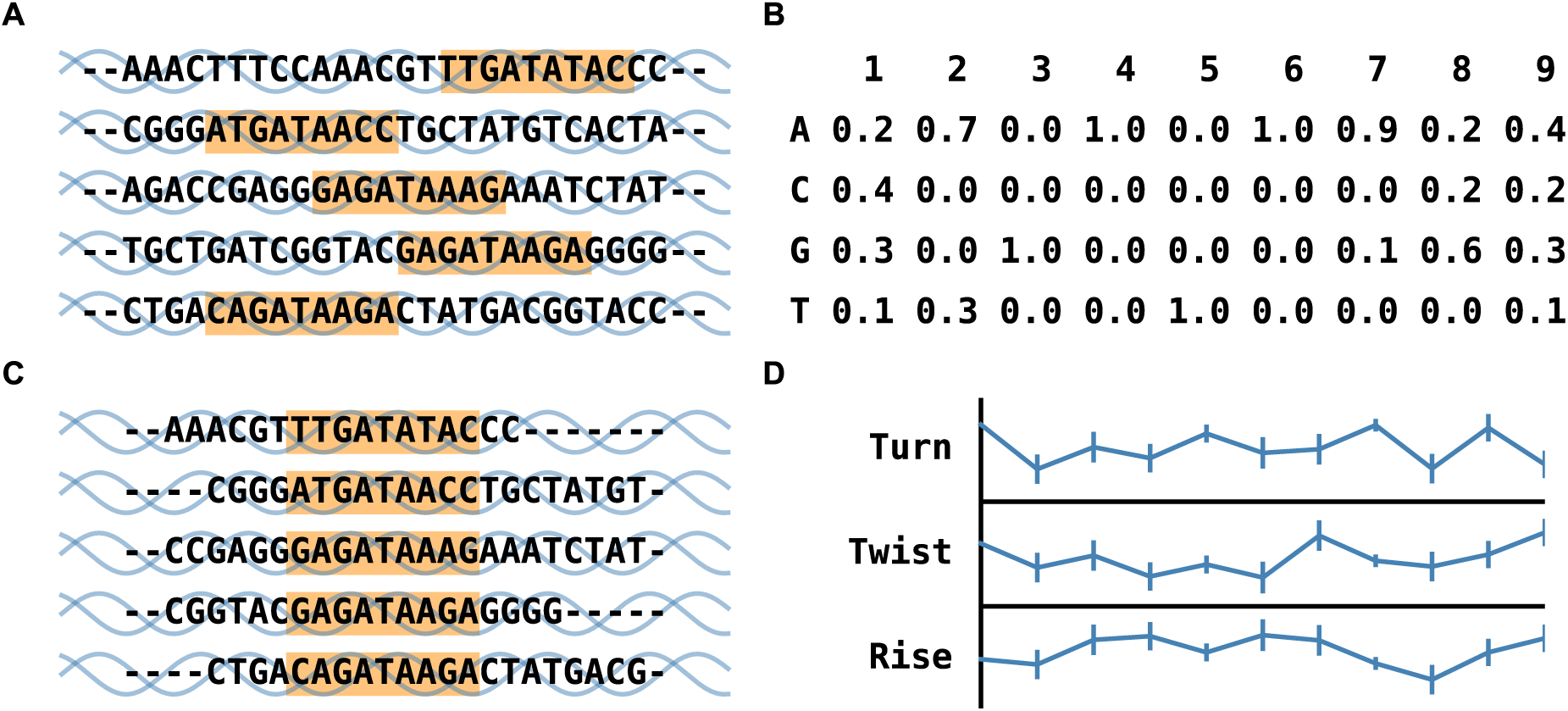
Graphical overview of structural motifs. **(a)** A series of DNA regions with a known binding site for GATA1 (orange). **(c)** These same regions are aligned by their binding site. **(b)** The alignment is used to calculate a distribution of base frequencies at each position, represented as a traditional PWM. **(d)** For the StruM paradigm, a distribution (mean, standard deviation) is computed at each position of the binding site for several shape features.

While most binding site representations are sequence-based, the physical interaction between the TF and TFBS must be compatible in both terms of electrostatics and sterics [26]. It has been shown that some transcription factors prefer specific shape configurations of the DNA [13, 27, 14]. It may be that the PWM and other sequence-based representations of TF binding motifs are abstracting these shape preferences, as sequence and DNA shape are tightly linked. We hypothesized that binding motifs could be modeled directly by DNA shape parameters.

The StruM model operates under the same basic assumptions as the PWM, but extends the model to correspond to shape values. If quantitative values can be obtained for characteristics of the DNA such as the Rise, Twist, and Turn, a distribution can be computed at each position of the binding site for these features. The StruM model parameterizes these distributions with the mean value and the observed standard deviation (Figure 1d).

Figure 2 shows the motif for FOXA1, learned using MEME [24]. Figure 2a is the traditional web logo representation of the PWM. A StruM trained on the same binding site regions identified by MEME show several interesting features (Figure 2b). The first thing to note is that there are indeed clear patterns. If there was no preference for a given shape, the average values would all be near zero, and the standard deviations would consistently be near one, given the scaled shape values used in the model. Rather, clear patterns are observed in both the average values and the variance. For example, the variance trends from high to low from left to right, corresponding to the information content in the PWM across the positions.

**Figure 2:**
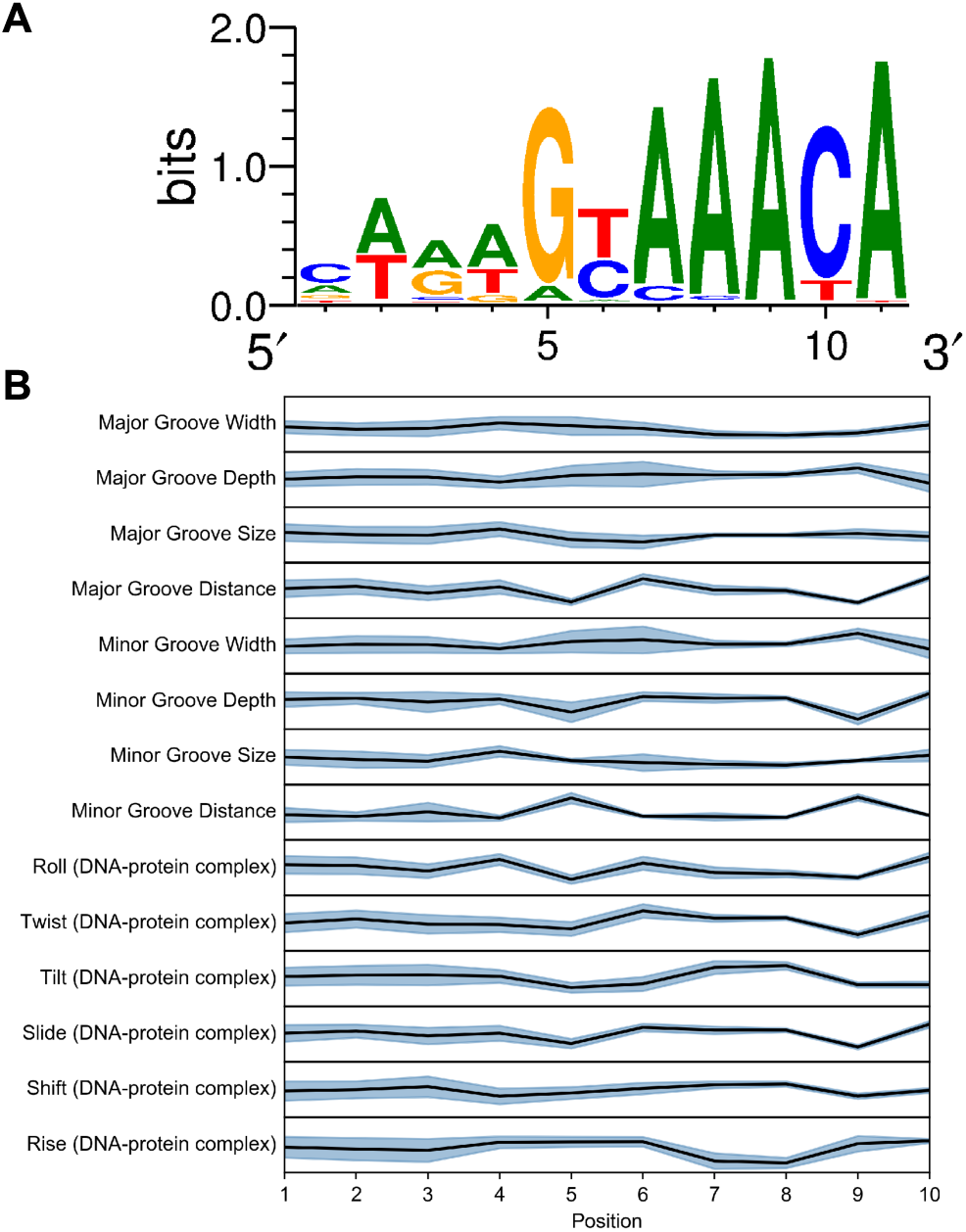
Motif for FOXA1. **(a)** Standard web logo representation of PWM for FOXA1. **(b)** Graphical representation of a StruM based on the same sequences as (a). The line plot represents the average value, and the shaded region is one standard deviation above and below.

### 3.2 StruMs specifically model TF binding sites

There is evidence to suggest that promoters and other similar genomic elements may share certain general shape features [28]. A simple example of conserved promoter structure would be the prevalence of the TATA-box [29, 30]. One possibility in evaluating the performance of shape based models to recognize TF binding sites is that they may be instead modeling general features of the type of genomic element that TF may target.

To determine whether a StruM is specific to the binding sites of the TF targeted in the ChIP experiment used to train the TFBS model the StruM was presented with a simple classification task to evaluate its ability to discriminate between ChIP peaks for its cognate TF from ChIP peaks deriving from other TF families. As a control, it was assessed whether StruM could distinguish between ChIP peaks from the same TF structural family versus the set of peaks from other TF families.

As shown in Figure 3a, StruMs were well able to distinguish between cognate TF binding sites and those belonging to other TF families (Avg. auROC = 0.77, Avg. auPRC = 0.77). In contrast, the control sequences appeared indistinguishable from the extra familial sequences (Avg. auROC = 0.53, Avg. auPRC = 0.54). Using a two-sided paired t-test this was a statistically significant difference (auROC: p-value = 4.33 × 10^−78^, auPRC: p-value = 1.62 × 10^−71^) indicating that the StruMs are specific to the TF on which they were trained.

**Figure 3:**
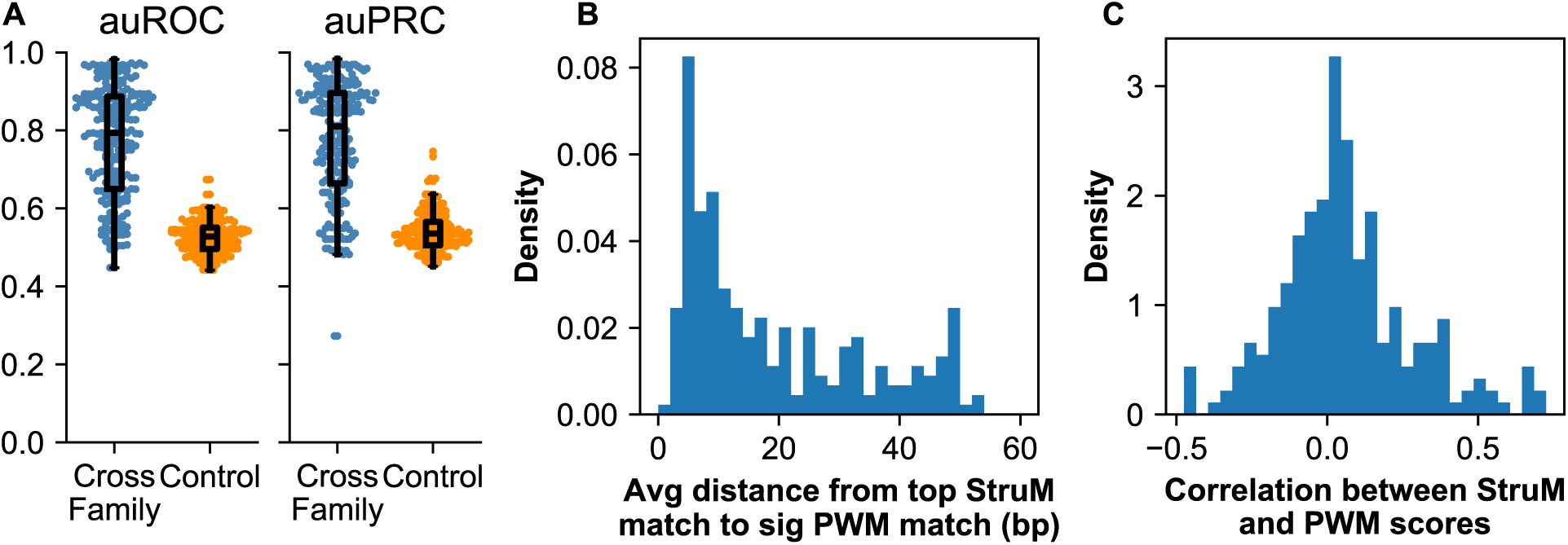
Specificity of StruMs. **(a)** The area under the ROC curve or PRC for TFs that could be unambiguously assigned to a TF family. Blue points correspond to the ability of the StruM to discriminate between ChIP-seq peaks for that TF *vs.* peaks for a different TF family (Cross Family). The orange colored points are the AUCs for the StruMs ability to discriminate between peaks from other TFs in the same family *vs.* peaks for a different TF family (Control). **(b)** Distribution of average distances from each StruM match to nearest significant PWM match. **(c)** The majority of correlations between scores assigned to kmers by PWMs and StruMs are near zero.

Once peaks are called from a ChIP-seq experiment and a motif is identified, the next step is frequently to identify the probable binding locations of the target factor at a higher resolution. One approach is to look for significant matches to the identified motif near the peak summit. If StruMs are accurately describing the binding sites of their cognate factors, high scoring positions within the peak should correspond with significant matches to the PWM. In order to evaluate this, FIMO [25] was used to identify significant matches to the PWM in the original ChIP sequences. The 200 bp surrounding each significant match was scored using the StruM, and the top scoring position for each sequence was retained. For each ChIP experiment, the average distance between each StruM match site and the nearest site identified by FIMO was calculated. As observed in the distribution in Figure 3b, the majority of StruMs recognized sites on average within 15 bp of the sites identified by FIMO. In many of these instances this offset seems to be the result of the motifs not aligning perfectly on the TFBS (Supplemental Figure S4, Supplemental Figure S8). The other peak in the distribution near 50 bp corresponds with StruMs that failed to accurately model the binding site (e.g. Supplemental Figure S4), as there is a strong negative correlation between the average distance and the auROC of the StruM model (Supplemental Figure S7, *R* = −0.69).

In the previous experiment, the PWM was used as the baseline for confirming the specificity of the StruM. To extend this analysis, a comparison was made of the ability of three models to discriminate between regions identified via ChIP-seq experiments and a negative control: either randomized peak sequences, or flanking sequences. The three models compared for each factor were: Position weight matrix (PWM), Dinucleotide weight matrix (DWM), and a Structural Motif (StruM). The area under the receiver operating characteristic curve (auROC) and the area under the precision-recall curve (auPRC) were calculated for each ChIP-seq experiment, using each of the models (Figure 4).

**Figure 4:**
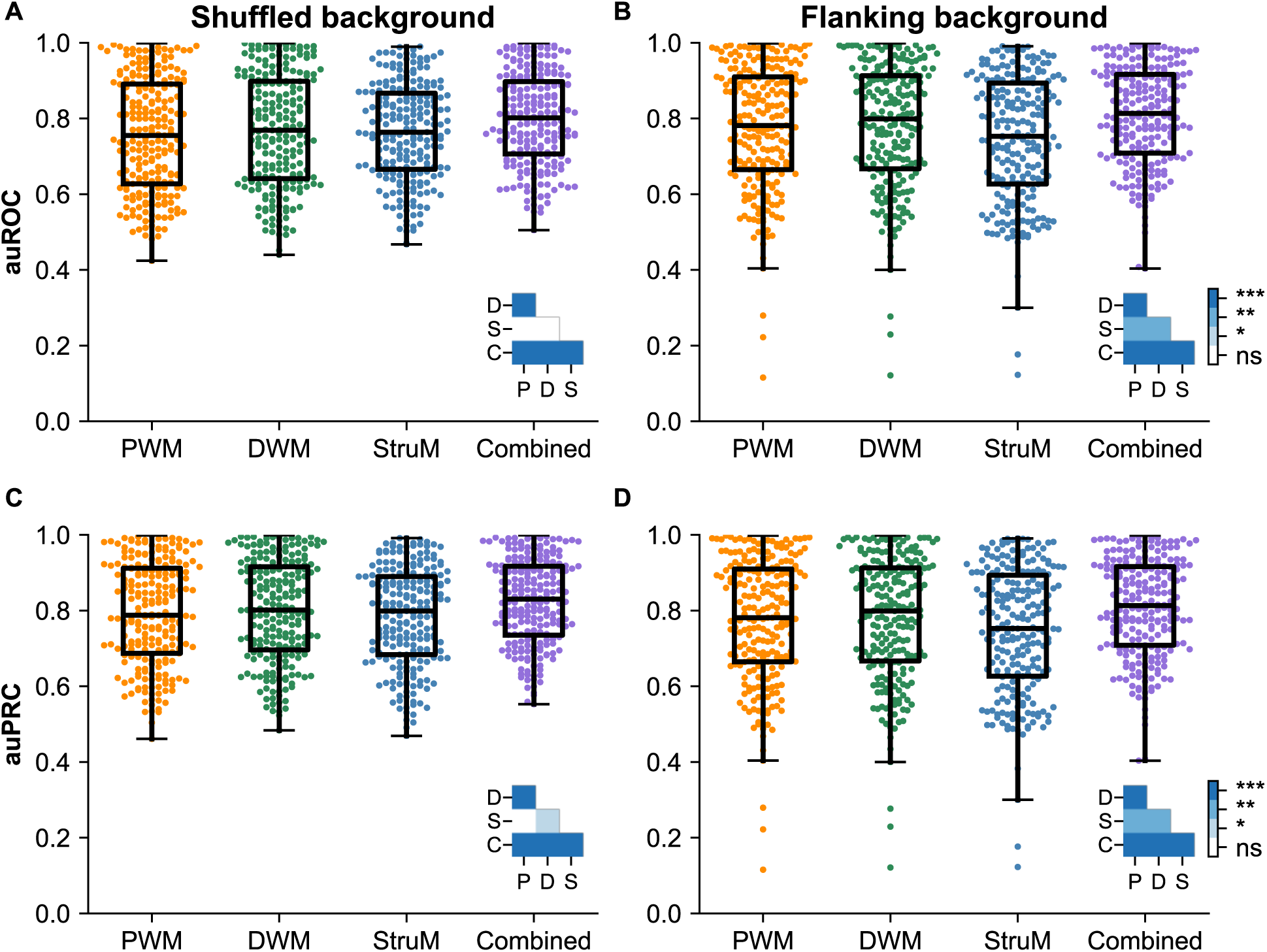
Classification of ChIP-seq peaks *vs* non-peak sequences. **(a-d)** Area under the curve for each TFBS model trained on the top 500 peaks, and tested using the next 500 sequences as the positive examples. The combined model used 10-fold cross validation to generate the score. The negative sequences came either from (a,c) shuffled peak sequences or (b,d) flanking sequences. Panels (a-b) represent the area under the ROC curve, and panels (c-d) represent the area under the PRC. Significance for each pairwise comparison is shown in the inset, with the following abbreviations: P–PWM, D–DWM, S–StruM, C–Combined model.

After evaluating 229 ChIP-seq experiments in K562 cells from the ENCODE consortium, StruMs perform at a similar level with the sequence based methods. It is interesting to note that using shuffled sequences as the negative set resulted in performance that was quite comparable across the three models. When using flanking sequences, the sequence-based approaches showed a slight but statistically significant edge over the StruM.

### 3.3 StruMs encode motifs differently than sequence-based methods

It is quite interesting to note that models constructed in this way (a probability distribution at each position of the motif, assuming independence between positions) perform relatively similarly, regardless of the feature being considered, be it mono- or dinucleotides, or shape features estimated from dinucleotides. In addition to similar performance in a simple classification problem, the different representations identify similar positions as being the putative targets for a given TF.

One might therefore expect there to be a strong correlation between the scores produced by each model for a given sequence. However when scoring a set of 1000 randomly generated sequences the models have near zero correlation (Figure 3c. Average correlation = 0.059, standard deviation = 0.22). This indicates that while the separate motif representations model the same site, they are encoding the information at that site very differently.

### 3.4 Shape and sequence are complementary

Given that both sequence- and shape-based methods can model the same sites with a similar level of accuracy, we investigated whether these methods were in fact redundant, despite showing very little relatedness between the ordering of random sequences. If these models are in fact parameterizing distinct features of the binding site, one would expect a combined model to outperform either model alone.

Towards this end a simple logistic regression model was trained for each TF, passing the maximum score from the PWM and the StruM as a length 2 vector for each sequence as input. Using 10-fold cross validation the combined model significantly outperformed each motif alone. (Figure 4a-d. Paired t-test; p-values < 4 × 10^−12^).

As further evidence of the complementarity of the models, the increase in auROC vs. shuffled sequences was plotted by the combined model over the StruM against the increase over the PWM performance (Figure 5a). In the event where the combined model simply agreed with the best performing model one would expect the points to fall along the x- and y-axes. Rather it was found that the majority of the points (68%) fall in the first quadrant off of the axes again indicating the complementarity of sequence and structural motif representations. Most of the remainder, accounting for 22% of all experiments, performed best with the StruMs alone.

**Figure 5:**
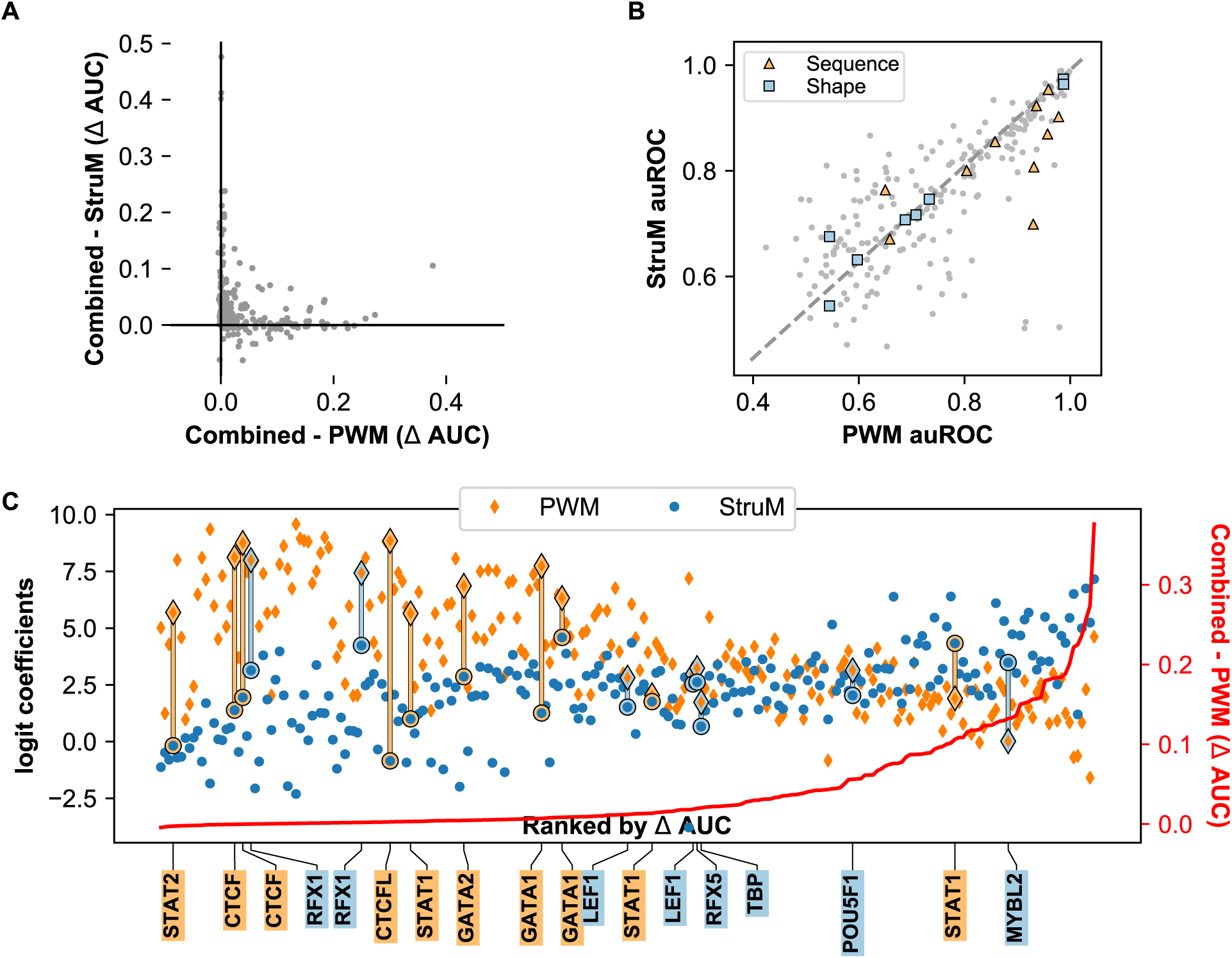
**(a)** The improvement of the combined model over StruMs (y-axis) was plotted against the improvement over PWMs (x-axis) for each TF. Points falling into the first quadrant show a positive impact on performance by joining multiple motif representations. **(b)** TFs that are known to utilize the base-readout mechanism (orange diamonds) fall on both sides of the line, while known shape-readers (blue circles) are generally better predicted by StruMs. **(c)** Coefficients for PWM and StruM from the combined logistic regression model were plotted against their rank improvement of the combined model over PWM alone. The absolute value of that improvement is indicated by the red line and corresponds to the values on the right-hand axis. The known sequence- and shape-readers are identified by gold and blue lines, respectively.

### 3.5 Towards distinguishing between direct- and indirect-readout mechanisms

Despite the similarity in performance on average between the sequence- and shape-based TFBS representations, there were several factors that showed a stronger than average preference for one method over another. We sought to understand whether these differences were limited to a subset of factors. First, the relationship between the AUC values for PWMs and StruMs for specific factors was examined (Figure 5b). Points falling along the dashed line representing *y* = *x* represent experiments where both representations perform equally well in predicting whether a sequence is likely to be bound by the factor.

We hypothesized that factors falling above the line employ shape- or indirect-readout mechanisms, whereas points below the line represent factors employing primarily base- or direct-readout. Several examples of known base- and shape-reading TFs are highlighted. In line with this hypothesis, those factors with a larger than average residuals tend to segregate by their readout mechanism.

Next, the coefficients of the combined logistic regression model were considered. One would expect that shape-readers would give more weight to the PWM than to the StruM. In direct contrast to the previous observations, the logistic regression model prefers the PWM a majority of the time, regardless of the annotated readout mechanism of the factor (Figure 5c). In fact only 1 out of 8 shape-readers weighted the StruM score more strongly, and the model preferred the StruM for only 1 out of 10 experiments for the known shape-readers.

## 4 DISCUSSION

In this work we have presented a novel representation of transcription factor binding site preferences termed **Stru**ctural **M**otifs, or **StruM**s. This model is an extension of the formulation behind the time-tested PWM that accommodates distributions of shape features. The flexibility of this model allows for variations of the StruM that can be tailored to specific tasks, and the incorporation and integration of additional data types. The DiProDB is used as a simple and fast system for converting sequences to a structural representation [16]. Other methods for shape estimation such as `DNAshapeR` package would fit well within this representation [31]. Arbitrary other data types such as DNase hypersensitivity may likewise be incorporated, as long as quantitative values can be generated for each position in the sequence. The StruM representation shows an ability to specifically model TF binding sites, as well as differentiate between ‘peak’ and ‘non-peak’ sequences in a ChIP-seq experiment.

Despite an average similarity to sequence methods in performance for these tasks, representing motifs using shape features is not universally appropriate. Some transcription factors employ a direct-readout mechanism whereby they recognize their binding site via interactions with specific base pairs. For these factors, the shape is an abstraction of the sequence information, rather than the other way around, and the PWM (or other sequence-based representation) is preferable for predicting binding sites.

A number of methods have been developed recently which seek to discover TF binding motifs in local DNA shape [32, 33]. As with StruMs, the values used for DNA shape are estimated directly from the sequence using a table like DiProDB [16] or from simulations like DNAshape [31]. The local structure of naked DNA is likewise determined by the sequence composition. This raises the question of whether consider higher order nucleotide features could fully capture the information contained in shape features. Indeed recent work has shown that given appropriate training data, dinucleotide features alone are sufficient to model most shape features [34, 11].

While it is possible to model the contribution of shape features indirectly through dinucleotide models, and indeed the dinucleotide model (DWM) displayed a strong performance, the StruM parameterizes the same information in a very different manner from these sequence-based methods. This disparity between the scores generated by the different model types turned out to be useful; combining the scores into a single model performed even better than either method alone. Thus the different mechanisms of encoding the same information are complementary to one another.

In summary, StruMs provide a novel way of considering transcription factor binding sites, which is complementary to sequence-based approaches. In fact many transcription factors may utilize a blend of direct- and indirect-readout mechanisms, agreeing with recent evidence (Reviewed in [13]). In this context, and given the observed complementarity, representing motifs using StruMs provides valuable information about the binding site preferences of transcription factors, and that when used in conjunction with sequence based methods can produce high confidence cell type-specific predictions of TFBSs. This will be valuable especially for studying the TF binding landscape of rare cell types for which carrying out extensive TF ChIP-seq experiments would be prohibited either by cost or the ability to collect enough cells for those experiments.

## 5 AVAILABILITY

Source code for StruMs and related tools as well documentation are available in the GitHub repository (https://github.com/pdeford/StructuralMotifs). All code required to replicate the analyses in this paper are available at (https://github.com/pdeford/strum_paper).

