## Supplemental Information for "DNA shape complements sequence-based representations of transcription factor binding sites"

#### 7 Supplementary Methods

##### 7.1 Training Structural Motifs

Two methods were applied to training the Structural Motifs used in this paper. These approaches differ in whether they take a sequence-centric approach to identifying the binding site. In the case where the training is directed by sequence, the kmers identified by MEME as contributing to the `meme-chip` PWM are taken as being representative of the binding site. Each one of these aligned binding site sequences is translated to 'structural space'. This is done by iteratively considering each dinucleotide in the sequence, and looking up the corresponding values in DiProDB for the features associated with the '`full`' filter mode in the StruM package (see Table 1). If the binding site identified by the PWM is of width $k$, and there are $p$ features, the size of the feature vector for this sequence is of length $(k-1) \cdot p$. The 'Maximum Likelihood' StruM, is then computed by taking the arithmetic mean and standard deviation at each of these $(k-1) \cdot p$ position-specific features across all of the aligned binding sites.

The second method used to train Structural Motifs was an Expectation-Maximization approach (algorithm described more fully below) directly on the structural representation of the training sequences. This version of the StruM was trained on the same set of sequences passed to MEME, i.e. the 100 bp surrounding the peak summit of the top 500 most enriched peaks in the ChIP experiment. In order to speed up the training time, the Expectation Maximization algorithm was done using the 'proteingroove' filter mode of the StruM package (see Table 2). This filter mode only uses 14 features related to the major and minor grooves and values derived from protein-DNA complexes. 10 random restarts were used to initialize the motif. After convergence of the models, the one with the highest likelihood was retained. This intermediate 'protein-groove' StruM was used to score the training sequences, and the best scoring kmer from each sequence was extracted. These were then translated to structural space with full 96 features available in the 'full' mode, and the parameters derived in a maximum likelihood fashion from this set of kmers.

| Count | Feature |
|---|---|
| 1 | Bend |
| 1 | Clash Strength |
| 2 | Enthalpy |
| 2 | Entropy |

| | |
|---|---|
| 1 | Flexibility_shift |
| 1 | Flexibility_slide |
| 9 | Free energy |
| 1 | Major Groove Depth |
| 1 | Major Groove Distance |
| 1 | Major Groove Size |
| 1 | Major Groove Width |
| 2 | Melting Temperature |
| 1 | Minor Groove Depth |
| 1 | Minor Groove Distance |
| 1 | Minor Groove Size |
| 1 | Minor Groove Width |
| 1 | Persistance Length |
| 1 | Probability contacting nucleosome core |
| 1 | Propeller Twist |
| 3 | Rise |
| 2 | Rise (DNA-protein complex) |
| 1 | Rise stiffness |
| 1 | Rise_rise |
| 4 | Roll |
| 2 | Roll (DNA-protein complex) |
| 1 | Roll stiffness |
| 1 | Roll_rise |
| 1 | Roll_roll |
| 1 | Roll_shift |
| 1 | Roll_slide |
| 2 | Shift |
| 2 | Shift (DNA-protein complex) |
| 1 | Shift stiffness |
| 1 | Shift_rise |
| 1 | Shift_shift |
| 1 | Shift_slide |
| 3 | Slide |
| 2 | Slide (DNA-protein complex) |
| 1 | Slide stiffness |
| 1 | Slide_rise |
| 1 | Slide_slide |
| 4 | Stacking energy |
| 3 | Tilt |
| 2 | Tilt (DNA-protein complex) |
| 1 | Tilt stiffness |
| 1 | Tilt_rise |
| 1 | Tilt_roll |

| | |
|---|---|
| 1 | Tilt_shift |
| 1 | Tilt_slide |
| 1 | Tilt_tilt |
| 1 | Tip |
| 6 | Twist |
| 2 | Twist (DNA-protein complex) |
| 1 | Twist stiffness |
| 1 | Twist_rise |
| 1 | Twist_roll |
| 1 | Twist_shift |
| 1 | Twist_slide |
| 1 | Twist_tilt |
| 1 | Twist_twist |
| 1 | Wedge |

Table 1: Features from the Dinucleotide Property Database used in the 'full' filtering mode of the StruM package. The 'Count' column represents how many times that feature appears, as the DiProDB table references some features multiple times, from different source in the literature.

| Feature |
|---|
| Major Groove Depth |
| Major Groove Distance |
| Major Groove Size |
| Major Groove Width |
| Minor Groove Depth |
| Minor Groove Distance |
| Minor Groove Size |
| Minor Groove Width |
| Rise (DNA-protein complex) |
| Roll (DNA-protein complex) |
| Shift (DNA-protein complex) |
| Slide (DNA-protein complex) |
| Tilt (DNA-protein complex) |
| Twist (DNA-protein complex) |

Table 2: Features from the Dinucleotide Property Database used in the 'proteingroove' filtering mode of the StruM package.

#### 7.2 Expectation Maximization training of StruMs

Our approach to expectation maximization was modeled after the OOPS model (only one per sequence) used by MEME [24]. Due to the formulation of StruMs as a combination of normal distributions, the parameters can be estimated using a variation of a weighted average.

##### 7.2.1 E-step

The likelihood ($l_{ij}$) of the $j$-th position in the $i$-th sequence being the start of the binding site is taken to be the score of the StruM at that position multiplied by the likelihood of the flanking regions matching the background model ($\phi_B$):

$$l_{ij} = \prod_{n=1}^{j-1} P(v_{ij}|\phi_B) \prod_{n=j}^{j+k-1} P(v_{ij}|\phi_{i-j+1}) \prod_{n=j+k}^{N} P(v_{ij}|\phi_B)$$

The likelihoods are then normalized on a by-sequence basis to produce $M$, the matrix of expected start positions:

$$M_{ij} = \frac{l_{ij}}{\sum_{j'=1}^{m} l_{ij'}}$$

##### 7.2.2 M-step

The maximization step takes these likelihoods and calculates maximum likelihood values for $\mu$ and $\sigma$ for each of the $m$ position-specific features:

$$\mu_j = \sum_{i=1}^{n} \sum_{\mathrm{v}} \frac{v_{ij} \cdot M_{ij}}{\sum_i \sum_j M_{ij}}$$

$$\sigma_j = \sum_{i=1}^{n} \sum_{\mathrm{v}} \frac{(v_{ij} - \mu_j)^2 \cdot M_{ij}}{\sum_i \sum_j M_{ij} - \frac{\sum_i \sum_j M_{ij}^2}{\sum_i \sum_j M_{ij}}}$$

#### 7.3 Filtering Position-Specific Features

As is the case with PWMs, not all position-specific features will contribute equally to the specificity of the motif. Analagously to positions with very low information content in a PWM, position-specific features in a StruM with large values for $\sigma$ don't reveal much information about the binding site, and can tolerate high amounts of variability at that site.

These non-specific features may contribute to the noisiness of the signal without appreciably contributing to the specificity of the motif. By filtering out non-specific features, not only might the signal to noise ration be improved, but also reduce the time required to score a kmer with the motif.

Two methods of identifying non-specific features in the StruM were explored. The first was simply based on the value of $\sigma$. Large values of $\sigma$ by definition correspond to large amounts of variation in the training data for that feature. Excluding position-specific features with a value for $\sigma$ greater than some threshold would limit the score for a sequence to only derive from specific features.

The second method considered was to use a Fisher score, based on the Fisher Linear Discriminant. This strategy requires a negative set, and for each feature compares the difference in mean values for the positive and negative sets, to the difference in variability observed for the positive and negative sets for that feature. More specifically, the Fisher score $V$ for the $i$-th position-specific feature can be computed as:

$$V_i = \frac{(\mu_{i+} - \mu_{i-})^2}{\sigma_{i+}^2 + \sigma_{i-}^2} \quad \text{(S1)}$$

A larger value for $V_i$ corresponds to a larger difference between the two sets, after accounting for their variability. After training the StruM, the positive set was constructed by taking the best scoring kmer from each training sequence. The negative set was then generated by randomly shuffling each of the kmers in the positive set.

In order to automatically identify a threshold for these two methods, the position specific features were rank-ordered by each of the metrics. A univariate spline was fit to these rank-ordered values. The point of inflection in this spline was selected as the threshold. Features with values above the threshold were retained for the $\log_{10}$Fisher score (Figure S1) and features with values below the threshold were retained for $\sigma$ (Figure S2).

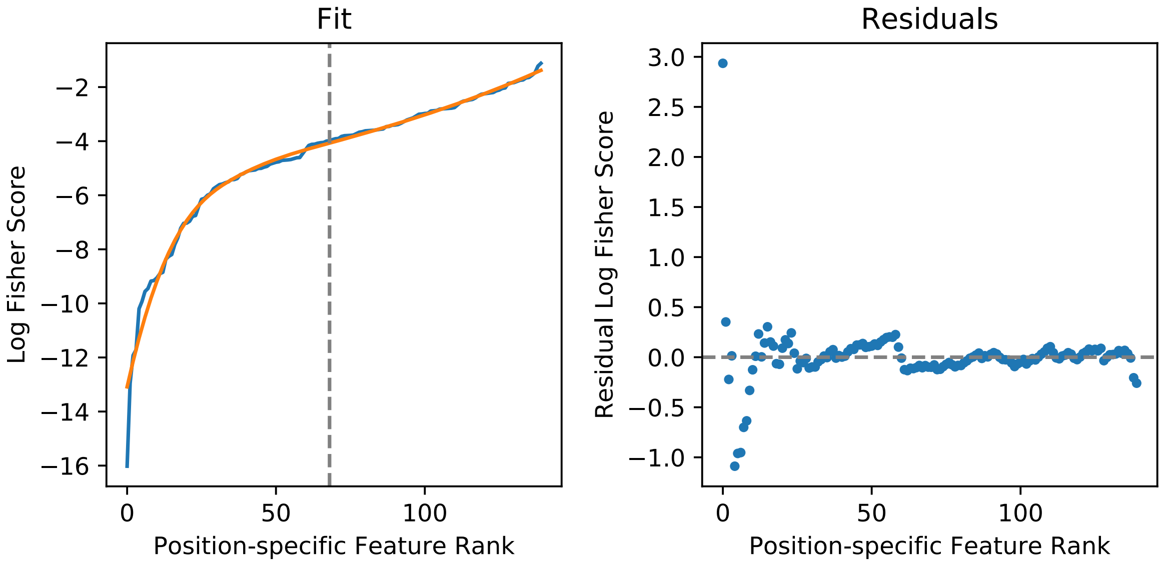


Figure S1: Position-specific StruM features were rank ordered by their log Fisher score determined from the binding sites, and a shuffled set of sequences. A univariate spline was fit (orange line) and the point of inflection determined as the threshold (vertical grey line). The residuals from the fit are on the right hand side.

The performance of using the full StruM and each of the filtered versions was evaluated using three metrics: The Fisher score, auROC, and auPRC (Figure S3). In general, the the filtered versions performed as well

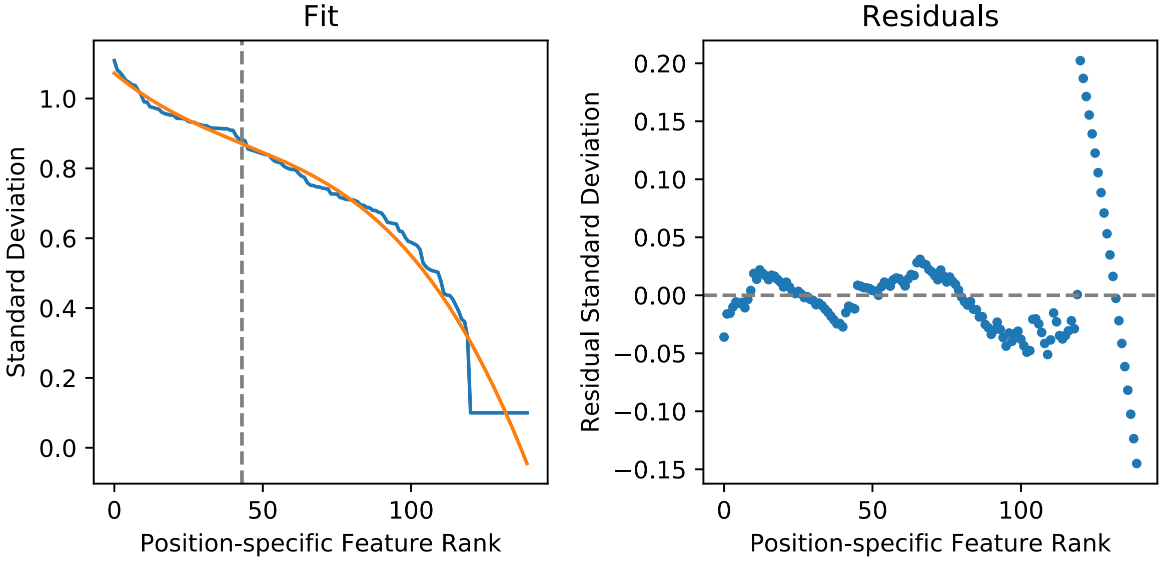


Figure S2: Position-specific StruM features were rank ordered by their value for $\sigma$. A univariate spline was fit (orange line) and the point of inflection determined as the threshold (vertical grey line). The residuals from the fit are on the right hand side.

as or slightly better than the original full version of the motif (data not shown). We therefore elected to use filtered-StruMs for this analysis. Specifically, filtering on the variance threshold was used as it does not require any sort of negative set to compute.

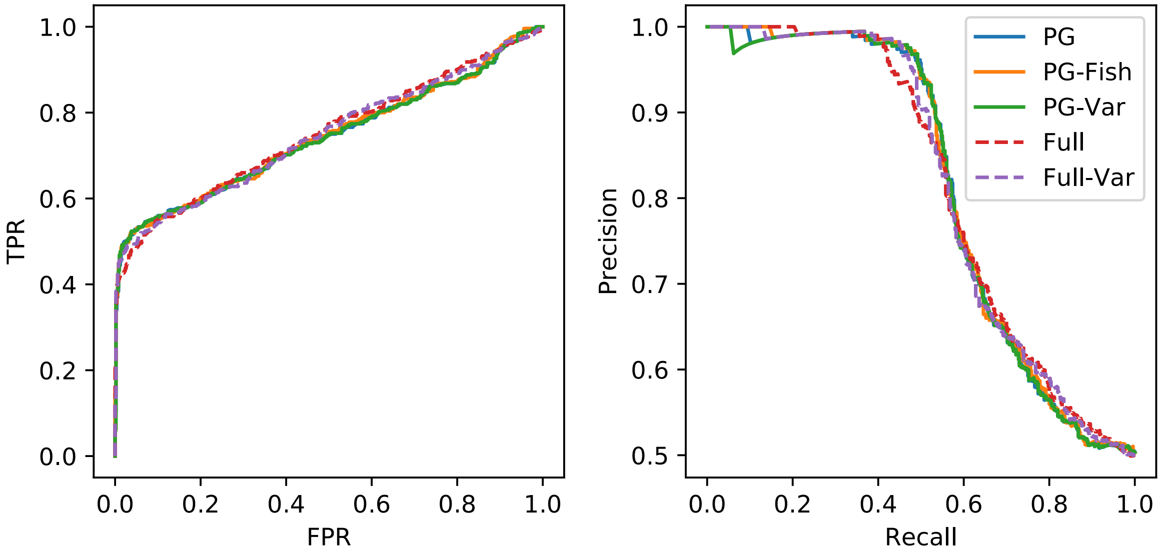


Figure S3: The performance of the the full StruM compared to the filtered version.

### 8 Supplementary Figures

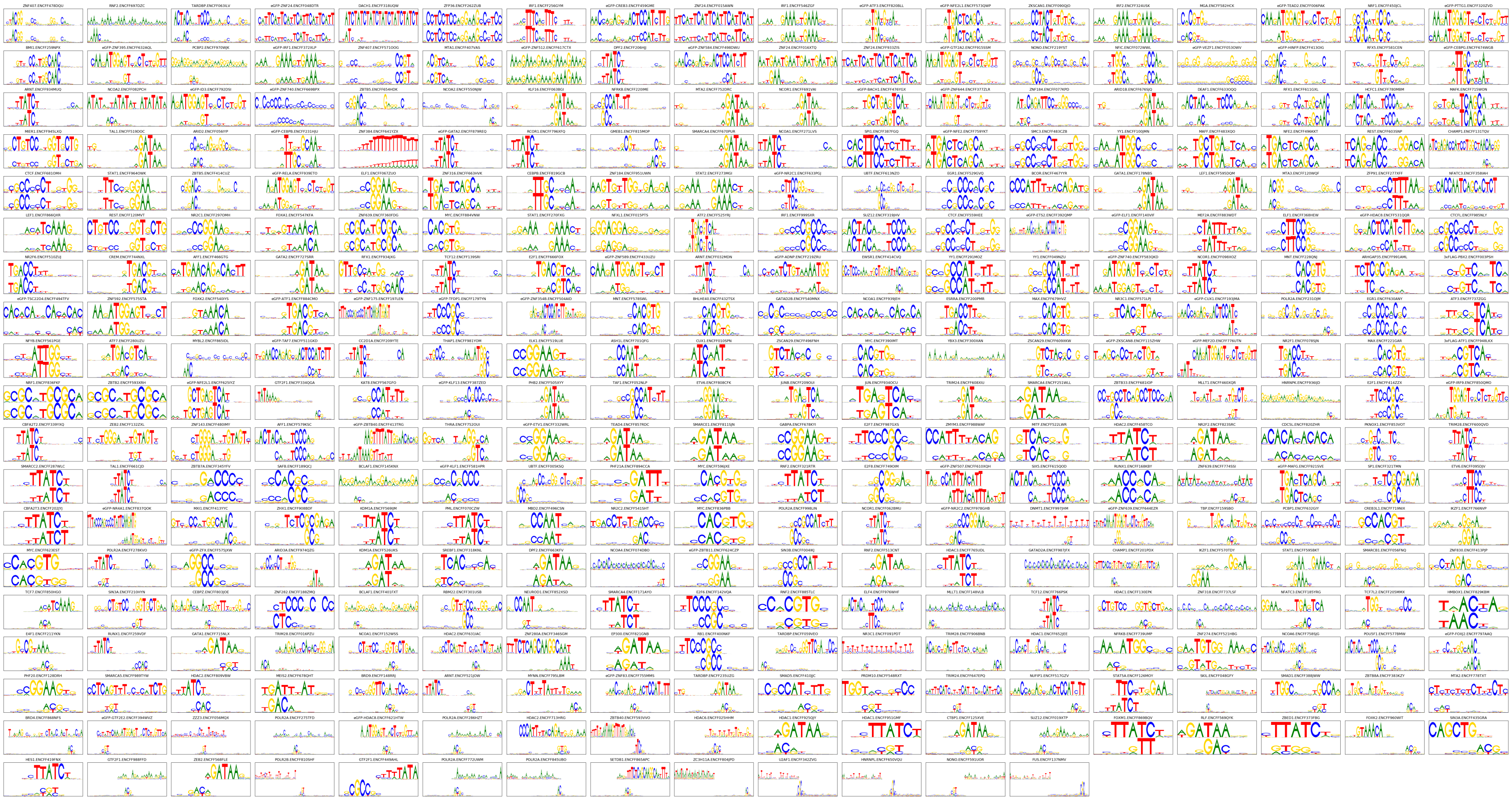

Figure S4: *MEME-trained PWMs aligned to motifs generated from StruMs*. As previously described, MEME was used to find a PWM for each ChIP experiment. Expectation maximization was concurrently used to generate a StruM for each experiment. In order to permit a more direct comparison of the motifs, the kmer in each training sequence with the highest StruM score was retained, and a PWM was generated from this set. The MEME-PWM and StruM-PWM were aligned by taking the retaining the maximum scoring alignment of either the forward or reverse complemented MEME-PWM to the StruM-PWM. The alignment score was calculated by summing across each aligning column in the motifs, where a column score is the Pearson correlation coefficient of the aligned columns. This figure shows the motif alignments sorted by their alignment score, with the highest scoring alignments at the top left. Rows were filled before columns.

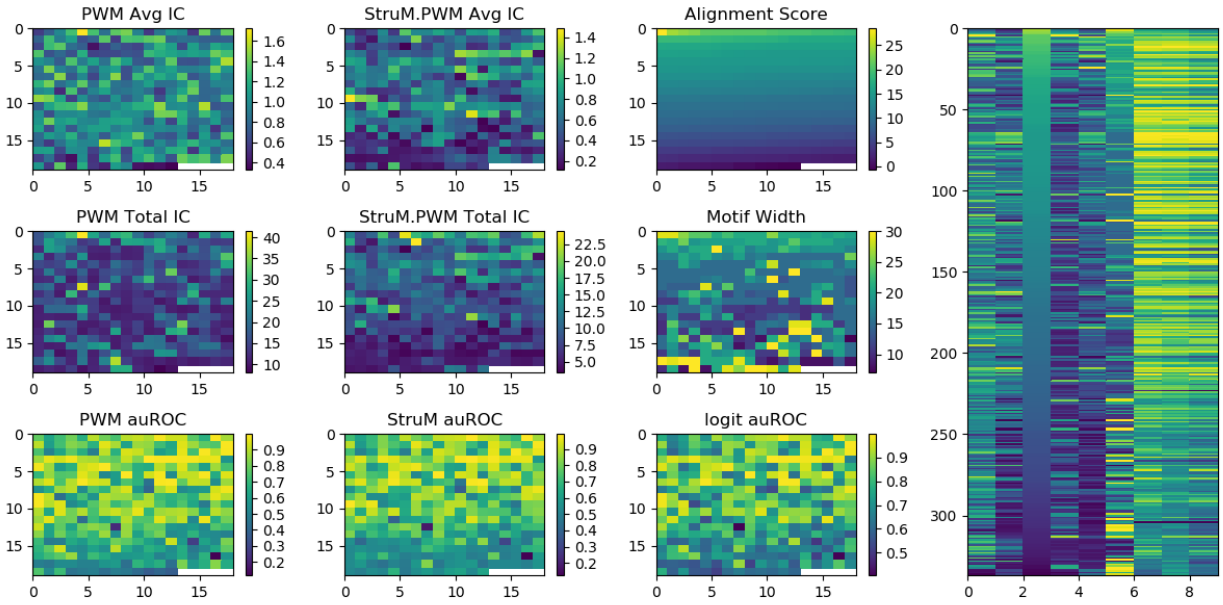


Figure S5: *Annotations for the motifs in Figure S4*. The cells in the heatmaps on the left side of this figure are organized to correspond directly with the position of the motifs in Figure S4. Several values are annotated. 'IC' is the Information Content of the motif, either summed across the full motif, or averaged. The alignment score is the summed Pearson correlation of the aligning columns in the aligned motifs. The motif width is the number of basepair positions represented in the motif. Finally, the auROC corresponds to the AUC using the shuffled sequences as background. The panel on the right hand side is the same nine sets of values, reorganized into a single normalized heatmap where each row represents the nine values for a single ChIP experiment.

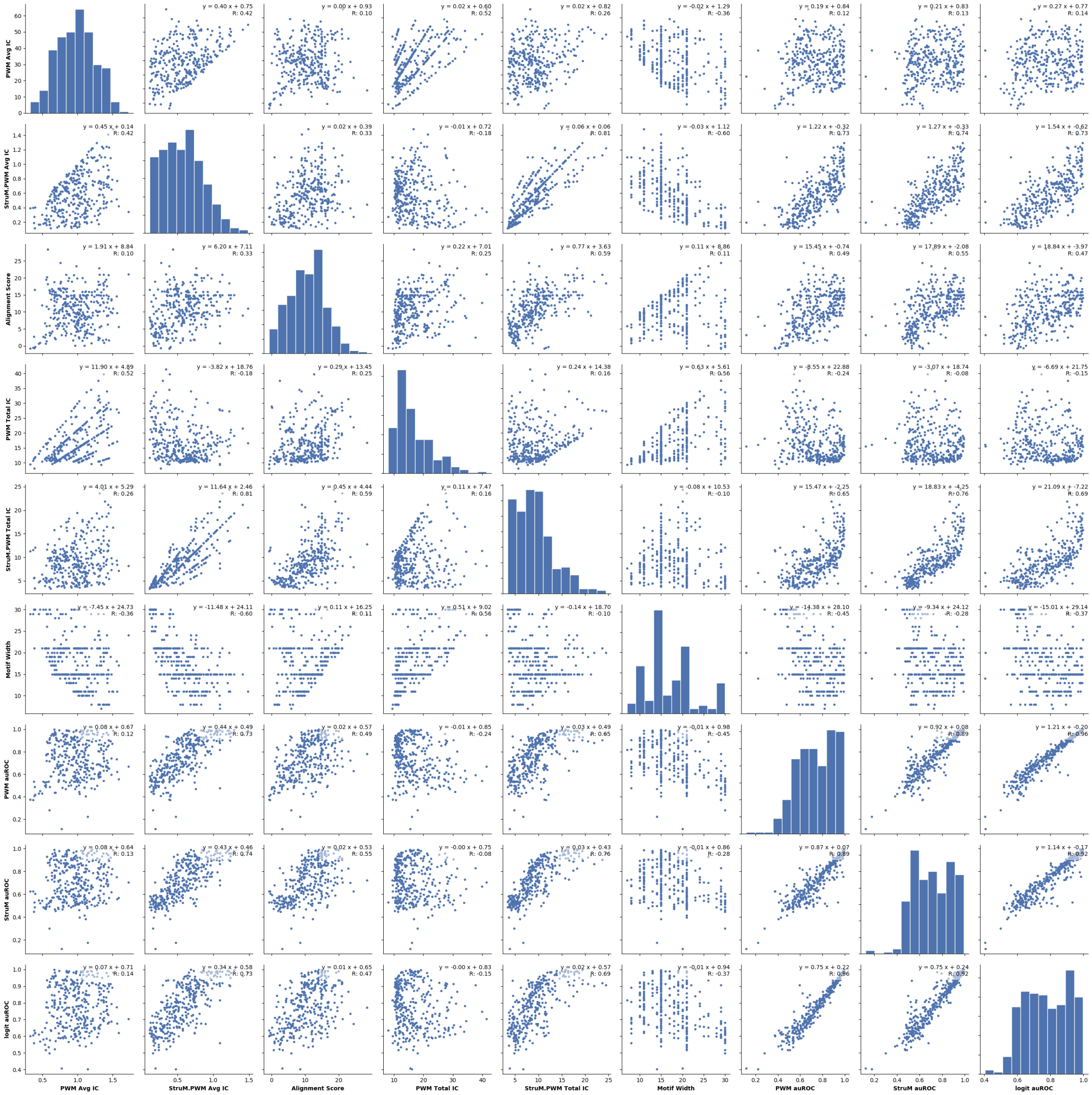


Figure S6: *Annotations for the motifs in Figure S4*. The values plotted here are the same as in Figure S5. In this case the pairwise relationships are shown between the separate variables. Each panel has the equation of the line of best fit as well as the correlation in the corner.

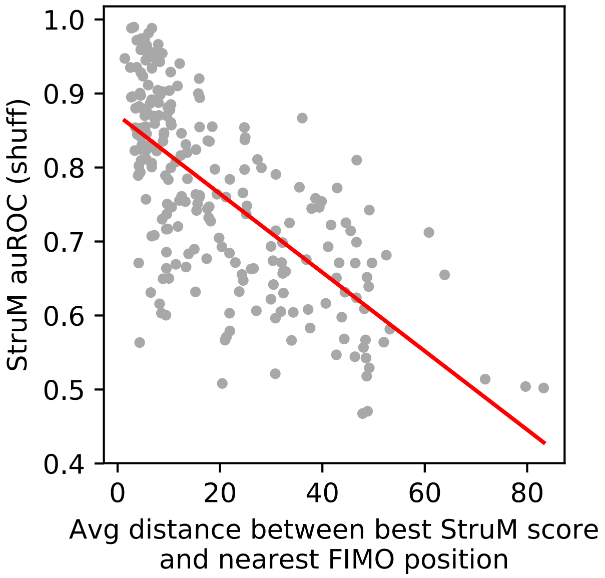


Figure S7: The performance of the StruM is inversely related to the distance between the top StruM matches and the nearest binding site identified by FIMO using the PWM.

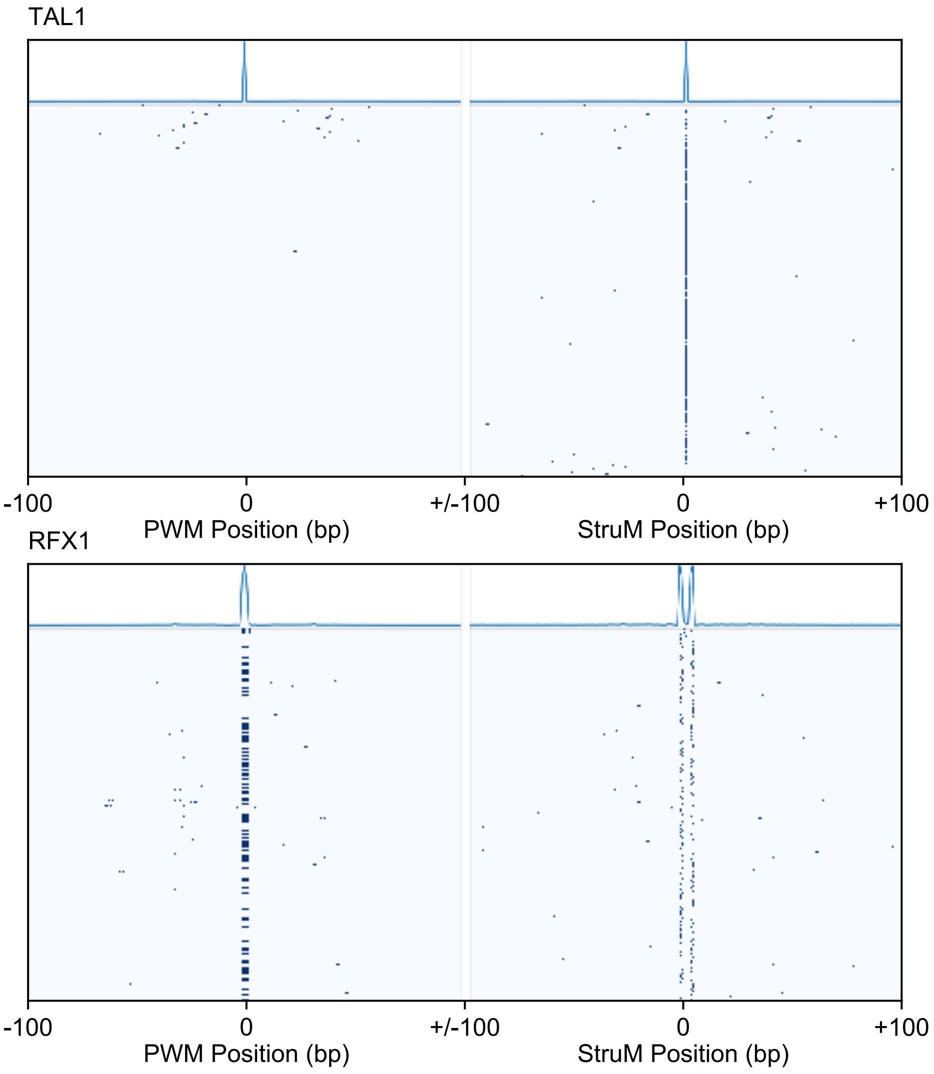


Figure S8: Distribution of top scoring StruM positions relative to PWM matches identified by FIMO. **(top)** Example of good correlation, small average distance with a single peak. **(bottom)** Example of good correlation, at a consistent small flanking distance..
